# Defining the transcriptional signature of esophageal-to-skin lineage conversion

**DOI:** 10.1101/2021.02.19.431899

**Authors:** Maria T. Bejar, Paula Jimenez-Gomez, Ilias Moutsopoulos, Bartomeu Colom, Seungmin Han, Fernando J Calero-Nieto, Berthold Göttgens, Irina Mohorianu, Benjamin D. Simons, Maria P. Alcolea

## Abstract

The ability of epithelial cells to rewire their cell fate program beyond their physiological repertoire has become a new paradigm in stem cell biology. This plasticity leaves behind the concept of strict stem cell hierarchies, opening up new exciting questions about its limits and underlying regulation. Here we developed a heterotypic 3D culture system to study the mechanisms modulating changes in the identity of adult esophageal epithelial cells. We demonstrate that, when exposed to the foreign stroma of adult skin, esophageal cells transition towards hair follicle identity and architecture. Heterotypic transplantation experiments recapitulated this cell fate conversion process *in vivo*. Single-cell RNA sequencing and histological analysis, capturing the temporality of this process, reveal that most esophageal cells switching towards skin identity remain in an intermediate state marked by a transient regenerative profile and a particularly strong hypoxic signature. Inhibition of HIF1a establishes the central role of this pathway in regulating epithelial cell plasticity, driving cells away from their transition state in favor of cell fate conversion.

## Main

Traditionally, epithelial tissues have been thought to be maintained and repaired solely by a pool of stem/progenitor cells (SCs) that balance proliferation and differentiation to preserve tissue integrity^1–7^. However, it has now become widely accepted that epithelial cell fate is more dynamic than originally thought^8^. Indeed, committed and differentiating cells retain the ability to reacquire developmental features and revert back to a stem cell-like state in response to tissue challenges^9–14^. However, the regulatory processes underlying cell fate plasticity remain poorly understood.

Under physiological conditions, epithelial plasticity now represents a well-established mechanism activated in response to tissue injury^15–17^. By rewiring their program of cell behavior, cells in the damaged area are allowed alternative cell fate choices, including the reactivation of their regenerative capacity^17–20^. This extends the pool of cells able to contribute to tissue damage beyond traditional SCs, guaranteeing a rapid and efficient healing process.

Further to this fate re-wiring in response stress, epithelial cells are able to change their identity well beyond their physiological constraints^21–25^. Tissue recombination experiments, dating back to the 1960s, established the ability of epithelial cells to change their identity and form appendages of a different nature, ranging from salivary and sweat glands to hair and feather formation, when exposed to the relevant ectopic stroma. It has been proposed that developing tissues respond more efficiently to these instructive signals^25,26^; however the response of adult tissues has been studied to a lesser extent. Despite efforts, the mechanisms that place epithelial cells in a permissive state, able to respond to foreign signals by changing their cell fate identity, remain to be elucidated. The inner workings dictating cell fate decision-making hold the key to identifying the principles underlying the full regenerative capacity of epithelial cells.

## Results

### 1. A 3D heterotypic culture system to study epithelial cell fate plasticity

In order to investigate the mechanisms governing cell fate plasticity, we developed an *ex vivo* model of esophageal-to-skin conversion, which comprises a change in cells identity between two architecturally similar squamous tissues of different embryonic origin^27^. The epithelium of the mouse esophagus (EE) and skin are both formed by layers of keratinocytes, supported by underlying stroma (dermis for skin, submucosa for esophagus) (**Fig.1A**). In both epithelia, proliferation is restricted to progenitors confined within the basal layer, which upon commitment to differentiation exit the cell cycle and stratify into the suprabasal layers^19,28^. The main architectural difference between the two tissues is that the mouse esophagus lacks any of the appendages typically found in the skin, including hair follicles (HFs), sebaceous glands and sweat glands^19^.

**Figure 1.**
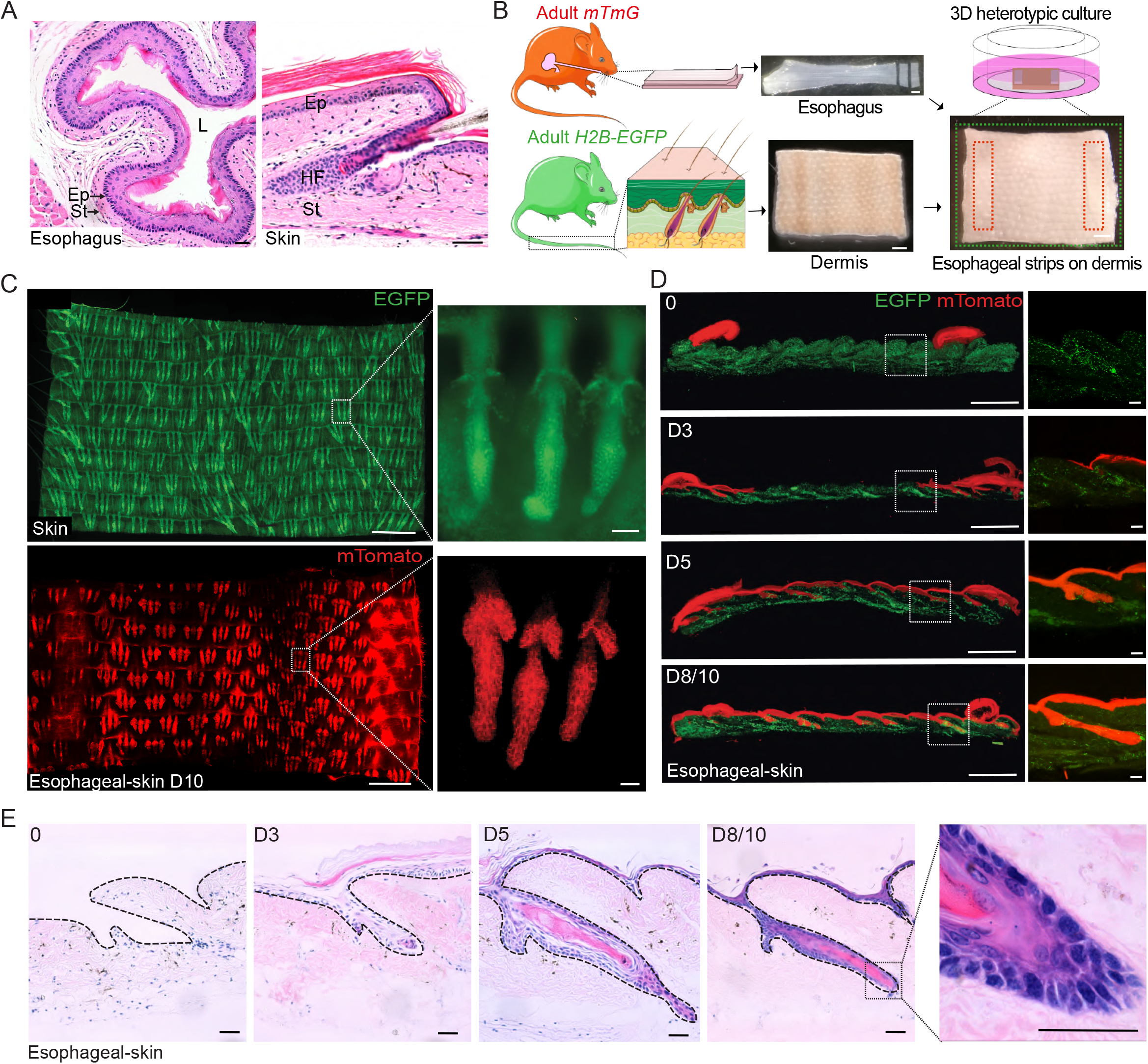
Esophagus-dermis 3D heterotypic culture as a model of ectopic niche regeneration. **(A)** H&E sections of adult esophagus (left) and skin (right) showing the characteristic tissue structure. Ep, epithelium; L, lumen; St, stroma; HF, hair follicle. **(B)** Schematic representation of the 3D heterotypic culture strategy. Esophageal epithelial strips obtained from adult tdTomato reporter mice were laid over denuded isolated dermis from H2B-EGFP mice and cultured for up to 10 days. During this time new epithelium forms between the strips, re-epithelializing the dermis. **(C)** Confocal images showing epithelial wholemounts of skin (top) from H2B-EGFP mice and esophageal-derived skin from heterotypic cultures (bottom) at 10 days (D10) as in **(B)**. Insets depict typical hair follicles (HF; top) and esophageal-derived HFs (bottom). **(D)** Cryosections of heterotypic cultures at the indicated time points throughout the 10 day culture. Insets show amplified images from the white-dashed squares, illustrating dermis re-epithelialization by the esophageal-derived cells (mTomato, red). n=3 animals. **(E)** H&E sections showing formation of esophageal-derived HFs in heterotypic cultures over a 10 day period. Dashed grey lines indicate basal membrane. **Scale bars.** 1A and 1E (50 μm); 1B-1D (1 mm), insets (50 μm). **Fluorescent reporters.** Green, EGFP; red, tdTomato.

For our *ex vivo* model we adapted an organotypic 3D culture system that we previously developed to study esophageal re-epithelialization during wound healing^19^. This system allows culturing combinations of epithelium and denuded stroma of different origins, in order to explore changes in cell behavior as the newly forming epithelium covers the stroma (**Fig.1B** **and SFig.1A**). Here, EE strips (5×1mm) were placed over a larger piece of denuded skin dermis (7×9mm) and the associated changes in epithelial cell identity during the re-epithelialization process were investigated (**Fig.1B**; **SFig.1A and B**). Of note, dermal papillae (DP), where the HF instructive signals are generated^21,29,30^, remained in the peeled dermis (**SFig.1C**). EE grown over esophageal stroma, was used as a control (**SFig.1A**).

In order to identify the origin of the cells generating the new structures in the heterotypic system, mouse strains constitutively expressing green or red fluorescent proteins were used. Specifically, EE was obtained from mTmG or nTnG mouse lines (constitutively expressing the red tdTomato reporter), while skin derivates came from H2B-EGFP mice (constitutively expressing a Histone-2B enhanced green fluorescent protein, EGFP), unless stated otherwise (**Fig.1A**). After growing the heterotypic cultures for 10 days (D) on minimal media, lacking any added growth factors (see methods), the red EE re-epithelialized the denuded green dermis and formed structures reminiscent of *in vivo* HFs (**Fig.1C**). The newly formed epithelium was sustained by basal cell proliferation, determined using the cell cycle reporter mouse line Fucci2a (**SFig.1D**), and did not present an increased Caspase3 activity compared to its *in vivo* counterparts (**SFig.1E**), confirming the integrity of the *in vitro* derived esophageal skin. By closely analyzing the temporality of the re-epithelization process, we observed that the first HF-like structures were formed by D3 post-culture (**Fig.1D** **and** **E**; **SFig.1F**). Full re-epithelialization was typically achieved at D5, but it was not until D8-10 that the newly formed epithelium thickened and keratinized (**Fig.1D** **and** **E**).

To measure the efficiency of esophageal-derived HF-like structures, we first quantified the number of HF units found in recipient pieces of Lgr5-CreGFP (green) skin prior epidermal peeling (**SFig.1G**). This was then compared to the number of *de novo* esophageal-derived HFs (red) formed over the same dermis at D10. The results estimate that 91.4±1.8% of HF “sockets” had been re-epithelialized with esophageal epithelium (**SFig.1G**).

### 2. Adult esophageal-to-skin lineage conversion in heterotypic organ cultures

We next set out to investigate whether our heterotypic culture system was enabling the EE to change its identity towards skin lineage.

To this end, wholemounts of the newly formed esophageal (e) derived epidermis (eSKIN) were immunostained for typical epidermal and esophageal markers, and analyzed by confocal imaging. The *ex vivo* skin structures of esophageal origin (eHF, esophageal-derived HF; eIFE, esophageal-derived interfollicular epidermis) were compared to i) *in vitro* skin (s) and esophageal controls generated as isotypic cultures (skin-on-dermis, sHF and sIFE; esophageal-on-submucosa, eEE), and ii) the respective *in vivo* tissue wholemounts (EE, skin HF and IFE). Most newly formed eHF showed a widespread expression pattern of the typical HF markers Keratin 17 (KRT17) and Keratin 24 (KRT24)^31–33^ (~78% and ~91%, respectively; **Fig.2A** **and** **B**). Reassuringly, expression of these proteins was present in HF controls, while not detected in the EE, eEE or eIFE controls (**Fig.2A**; **SFig.2A and B**). Conversely, expression of Keratin 4 (KRT4), a specific marker of differentiated esophageal cells, was downregulated in the eSKIN (**SFig.2C**), in line with previous observations^21^. These results indicate that esophageal cells exposed to the ectopic stroma of the dermis in our heterotypic cultures are transitioning towards skin identity.

**Figure 2.**
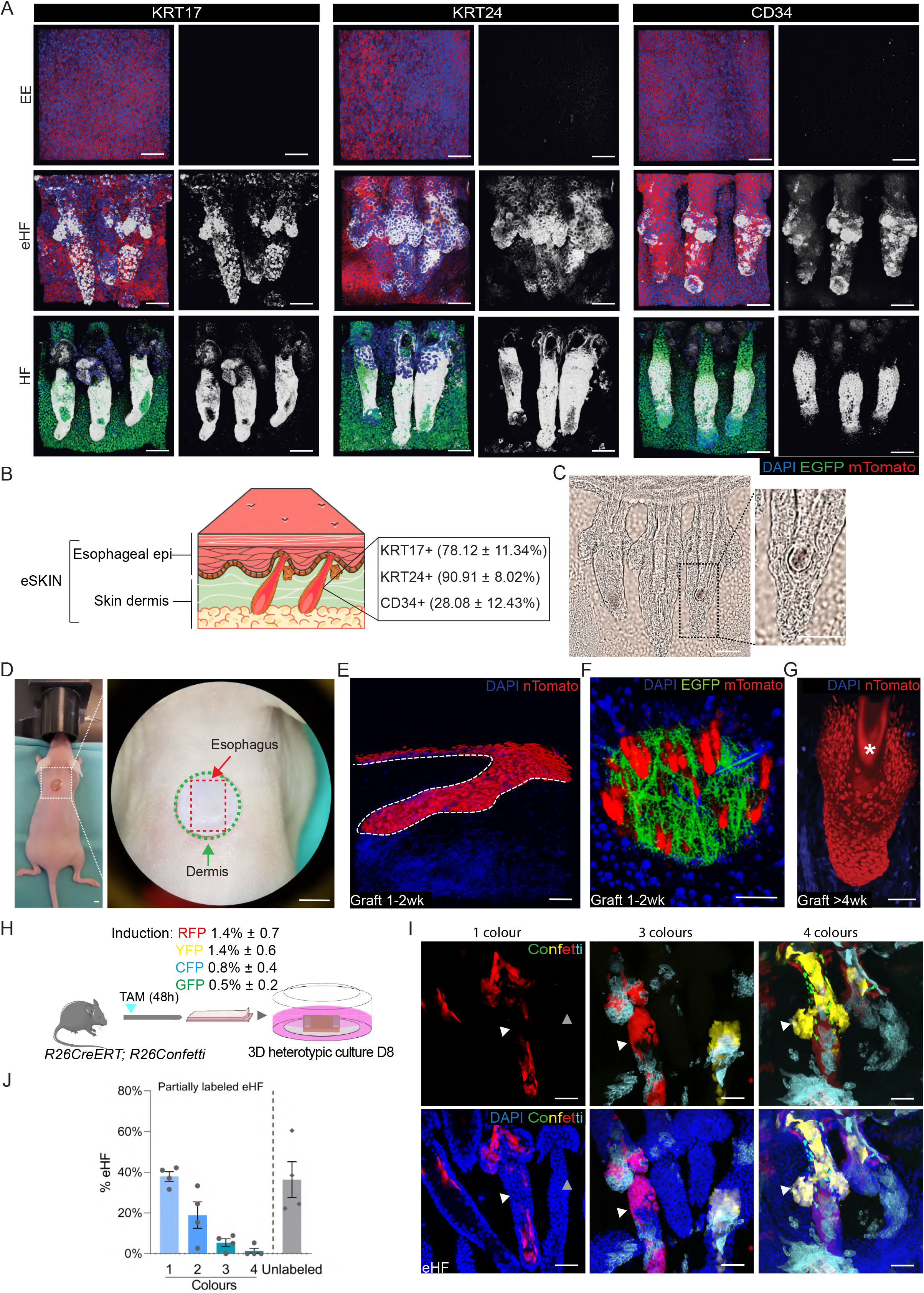
Esophageal cells undergo changes in cell identity when exposed to denuded dermal skin *in vitro* and *in vivo*. **(A)** Representative 3D rendered confocal z-stacks showing expression of the typical hair follicle (HF) markers KRT17 (left columns) and KRT24 (middle columns), as well as the HF stem cell marker CD34 (right columns). Esophageal epithelium (EE) from membrane tdTomato mice (EE, top panels), esophageal-derived HFs (eHF) from heterotypic cultures as in **Fig 1B** (middle panels), and skin HF from H2B-GFP mice (Bottom panels). For each marker, n=3 animals. **(B)** Illustration indicating the percentage of eHF units expressing KRT17, KRT24 and CD34, relative to total eHF, quantified from images in (**A**). Presented as mean ± SEM. n=3 animals. **(C)** Brightfield images of an eHF triplet. Inset shows pigmented structures in the eHF. **(D)** *In vivo* transplantation experiment. An 8 mm punch biopsy was taken from the back skin of recipient nude mice, immediately followed by esophageal-skin graft transplantation of red EE cells on green skin dermis (inset). n= 6 independent experiments. **(E)** Confocal 3D image showing a full-thickness graft section at 10 days upon transplant. **(F, G)** 3D rendered confocal images showing engrafted eHFs at 2 weeks post-transplant (**F**) or 4 weeks post-transplant (**G**). Asterisk in (**G**) indicates structure reminiscent of a hair shaft. **(H)** Experimental protocol. R26^CreERT^R26^Confetti^ mice were injected with tamoxifen (TAM) to induce Confetti labeling. 48 hours later, the esophagus was collected and cultured on top of wild-type dermis as 3D heterotypic cultures for 8 days. Confetti reporter induction efficiency is shown as the percentage of RFP, YFP, CFP or GFP positive cells relative to EE basal cells at 48 hours post-induction (mean ± SD, n=3 animals). **(I)** Confocal images illustrating several combinations of Confetti labeled areas in eHFs from (**H**). n=3 animals. Red, RFP; yellow, YFP; Cyan, CFP and green, GFP. **(J)** Percentage of eHFs labeled with one or more Confetti reporter proteins (RFP, YFP, CFP or GFP) or unlabeled, relative to total eHF units. Presented as mean ± SEM. n=3 animals. **Scale bars.** 2A, 2C, 2E, 2G and 2I (50 μm); 2D (4 mm) and 2F (1 mm). **Fluorescent reporters and stainings.** Unless otherwise stated green is EGFP (skin origin) and red is mTomato. White, HF markers KRT17 (2A, left columns), KRT24 (2A, middle columns) and CD34 (2A, right columns).

Interestingly, although most eHFs exhibit marked levels of KRT17 and KRT24 staining, only a subset of those (~28%) expressed CD34, a typical HF stem cell marker not expressed in the EE (**Fig.2A** **and** **B**; **SFig.2A**)^19^. Moreover, the number of CD34+ cells in eHFs was widely distributed, ranging from 1 to 59 cells per HF-like structure (with an average content of 6.79 ±2.21 cells per eHF). The heterogeneous induction of CD34 expression suggest a halted process; where the majority of eHF cells are able to initiate the identity switch, but subsequently appear to remain in an intermediate stage of the cell conversion process.

These results support the notion that our heterotypic culture system is favoring EE cell fate conversion towards skin lineages. This was further reinforced by the emergence of seemingly pigmented hair shaft-like structures in the newly formed eHFs (**Fig.2C**).

### 3. Adult esophageal-to-skin lineage conversion in vivo

To investigate whether the esophageal-to-skin cell fate conversion observed *in vitro* could be reproduced *in vivo*, we developed a heterotypic transplantation technique recreating our *in vitro* strategy.

For this, an area of ~50mm^2^ skin was excised from the back of nude mice, and replaced by peeled tail dermis (expressing EGFP) overlaid with a strip of EE (expressing tdTomato; **Fig.2D**). 1-2weeks following the procedure, EE cells had re-epithelialized the dermis, generating HF-like structures that expressed the HF marker KRT24, and contained a central fiber resembling the HF shaft (**Fig. 2E** **to** **G**; **SFig.2D**). Similar experiments transplanting wild-type dermis overlaid with strips of EE from the cell cycle reporter mouse line Fucci2a, indicated that the newly generated *in vivo* HFs were actively proliferating (**SFig. 2E**). This data confirms previous observations^21^, supporting the ability of EE to form viable HF structures *in vivo*.

Since our heterotypic *in* vivo technique had a limited grafting life of approximately 4 weeks, it was unable to render any outgrowing hair shaft. Hence, in order to test the hair forming ability of esophageal cells further, we performed a second *in vivo* experimental protocol^34–36^. Specifically, we transplanted subcutaneously a mix of dissociated cells from neonate skin (expressing EGFP) and adult EE cells (expressing tdTomato) into the back of nude mice. This protocol resulted in the emergence of esophageal/epidermal (tdTomato/EGFP) chimeric HFs that generated new functional hair shaft-like structures 6 weeks post-grafting (**SFig.2F**).

### 4. Widespread contribution of adult EE progenitor cells during tissue identity conversion

To understand the cellular dynamics leading to eHF formation in our *in vitro* model, we turned to genetic linage tracing. R26^CreERT^R26^Confetti^ mice were induced *in vivo* resulting in random cell labelling of cells with one of four fluorescent reporters (YFP, CFP, RFP or GFP)^37^. 48 hours post-induction, the EE presented a clonal recombination efficiency that ranged between ~1-3 labelled cells per 200 basal cells (**Fig.2H**). At this time point, EE were collected and grown over wild-type denuded dermis for 8 days, following our heterotypic approach detailed above. *In vitro* tissues were peeled and Confetti reporter labeling quantified by confocal microscopy to track the behavior of the cells forming the eSKIN (**Fig.2I**). The analysis shows that, out of the ~4.5% originally labelled epithelium, ~63% of the eHFs presented labeling, a percentage much higher than expected if only a portion of the EE cells contributed towards the eHF formation (**Fig.2H** **and** **J**). Hence, these results suggest that EE cells undergo a process of clonal expansion and contribute in a widespread manner to *de novo* HF formation, in line with previous observations in EE wound healing^19^.

When exploring the reporter distribution within the eHF, no monoclonal events were found, i.e. no eHF was found to be fully colonized by cells marker by a single reporter. Instead, all labelled eHFs were incompletely labelled by one or more of the four Confetti reporters (**Fig.2I** **and** **J**). The majority were partially labelled by one (~38%) or two (~19%) reporters, with only a small percentage (~6.7) showing more than three colors. Together, this data reveals the polyclonal origin of the newly formed eHFs.

### 5. Single cell RNA sequencing analysis captures the regenerative signature of esophageal-derived skin

We next investigated the molecular mechanisms regulating epithelial cell plasticity and cell fate conversion in our heterotypic culture system using large-scale transcriptomics at single cell resolution.

To resolve the temporality of the esophageal-to-skin identity switch, we performed single-cell RNA sequencing (scRNA-seq) at two time points spanning the re-epithelialization process (D3 and D10) (**Fig.3A**); with D3 representing an early regenerative stage, and D10 the steady state reached at the experimental end-point. Two different sample types were analyzed for each time point: EE cells growing over their own stroma (eEE-D3, eEE-D10) and EE growing over the ectopic stroma of the skin (eSKIN-D3, eSKIN-D10). Sequenced samples contained viable tdTomato EE-derived cells (from nTnG mice), spiked with 5% EGFP epidermal host cells (sSKIN), as a reference control for cell identity (**Fig.3A**; **SFig.3A**).

**Figure 3.**
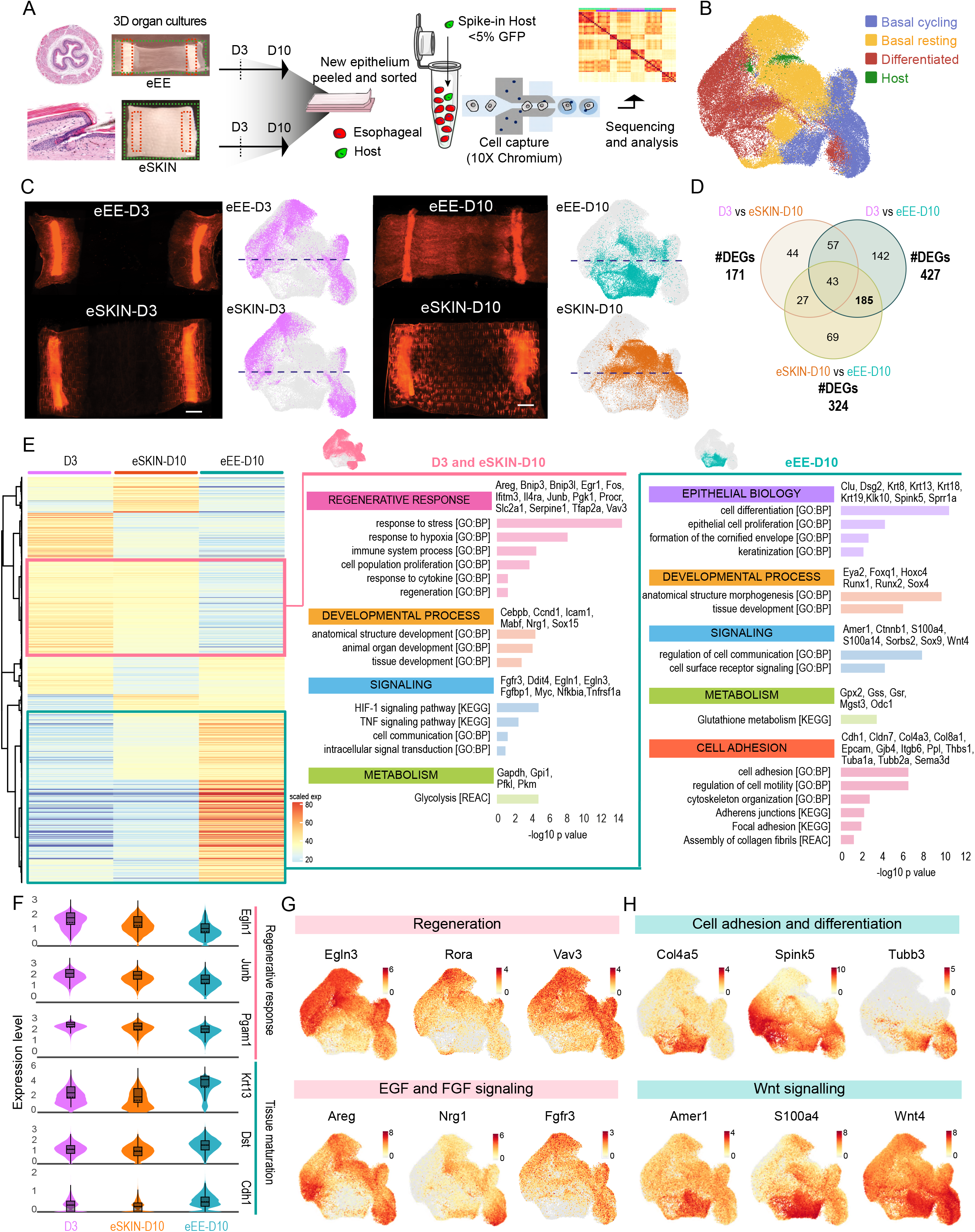
Single-cell transcriptional analysis shows active regenerative features are retained in esophageal-derived skin. **(A)** Overview of the single-cell RNA-seq experimental workflow (10x Genomics platform) for tdTomato^+^ eEE and eSKIN sorted cells, and spiked EGFP^+^ host-derived epidermal cells. **(B)** UMAP representing annotated cell types. **(C)** Representative images of eEE and eSKIN heterotypic cultures at days 3 (D3) and 10 (D10) post-culture shown adjacent to their respective UMAPs cell distribution. **(D)** Venn diagram showing the overlap of differentially expressed genes (DEGs) in the indicated comparisons across the three transcriptional branches: D3 (eEE + eSKIN), eSKIN-D10 and eEE-D10. Only basal cells were considered in the analysis. **(E)** Heatmap of DEGs between at least two of the three main branches (D3, eEE-D10 and eSKIN-D10) generated on scaled expression levels. Tables on the right show enriched signatures for the groups of genes up-regulated in D3/eSKIN-D10 (pink) and eEE-D10 (cyan), compiled from Gene Ontology Biological Process (GO:BP), Kyoto Encyclopedia of Genes and Genomes (KEGG) and Reactome (REAC) databases, depicting p-values and representative genes for each signature. **(F)** Violin plots illustrating the distribution of expression of genes associated with regeneration and tissue maturation for the three transcriptional branches (D3, eSKIN-D10 and eEE-D10). **(G,H)** UMAPs showing expression of genes up-regulated in D3/eSKIN-D10 (**G**, pink) and eEE-D10 (**H**, cyan) from the enriched signatures in **(E)**.

The combined scRNA-seq dataset was mapped using principle component analysis based on UMAP projection, followed by unsupervised clustering, and annotation, using signatures of known lineage and cell cycle markers (**Fig.3B**; **SFig.3B, H and I**; see Methods). Consistent with *in vivo* observations^19,38^, we identified three distinct epithelial cell populations for all sample types: i) actively cycling basal progenitors; ii) resting/committed basal cells; and iii) differentiated cells (**Fig.3B**; **SFig.3H and I**). We found the epidermal spiked-in population restricted to an individual cluster (cluster 18, Host). This was evidenced by detection of genes known to specifically discriminate EGFP host cells *versus* tdTomato esophageal cells (*Egfp* and *Gt(ROSA)26Sor*; **SFig.3C and D**). Note, *Gt(ROSA)26Sor* expression is blocked in nTnG esophageal cells due to the nTnG construct targeting into the Rosa26 locus^39^. Gene classifiers, based on a semi-supervised model that extends transcriptomic signatures of the detected tdTomato and EGFP cells to unlabeled cells, provided further reassurance that C18 represented a cluster significantly enriched in EGFP Host cells (**SFig.3E**). Expression of *Sox2*, a known marker of the basal layer of the adult EE^40,41^ found to specifically label cells of esophageal origin *in vitro* (eSKIN, **SFig.3F**), was largely absent in C18 (**SFig.3G**). This data confirms that C18 is enriched for cells of non-esophageal origin, and supports its Host epidermal identity.

The UMAP distribution of the single-cell data revealed a noticeable overlap between D3 esophageal cells growing over their endogenous stroma (eEE) and those growing over the ectopic dermis (eSKIN; **Fig.3C**). This suggested that esophageal cells at early re-epithelialization stages have a similar transcriptional signature, independent on their substrate. Hence, D3 samples were grouped when exploring gene expression patterns. The transcriptional profile of both eEE and eSKIN showed significant changes from D3 to D10, following tissue re-epithelialization, evidenced by a progressive shift from the top to the bottom of the UMAP space. Interestingly, the two D10 samples did not overlap; eSKIN retained an intermediate distribution between eEE-D3 and eEE-D10 cells grown on their native stroma (**Fig.3C**).

Next, to shed light on the regulatory processes governing the epithelial cell fate transition, we focused our analysis on the basal cell compartment, where progenitor cells reside^19^. Remarkably, when analyzing the 3 major transcriptional branches (D3, eSKIN-D10 and eEE-D10), we found that D3 and eSKIN-D10 presented a closer molecular signature, showing less differentially expressed genes (DEGs; 171 D3 vs eSKIN-D10) relative to other comparisons (427 eEE-D10 vs D3, and 324 eEE-D10 vs eSKIN-10D; **Fig.3D** **and** **E**). Esophageal control eEE-10D had the most distinctive transcriptional profile of the three main branches. This hinted at D3 and eSKIN-D10 cells having a closer resemblance at the transcriptional level. Gene Ontology (GO) enrichment analysis of DEGs revealed a particularly strong regenerative signature shared between D3 and eSKIN-D10 cells (**Fig.3E** **to** **G**). Indeed, pathways involved in active wound healing and tissue repair were found to be enriched in both branches; these included EGF (*Areg*, *Nrg1*, *Egr1*, *Vav3*, *Tfap2a*)^42–45^, AP-1 (*Junb*, *Fos*)^11,46,47^, and FGF (*Fgfr3*, *Fgfbp1*)^48^. Epithelial mesenchymal transition (EMT) related genes, known to be modulated during epithelial regeneration, showed a similar trend. Among these, we found targets downstream of HIF1a (*Pgk1*, *Slc2a1*, *Egln1*, *Egln3*, *Pfkl*, *Rora*)^49^, as well as *Igfbp2^50^*. Accordingly, D3 and eSKIN-D10 cells also showed an active stress response; displaying increased levels of genes promoting cell migration (*Bnip3*, *Bnip3l, Procr, Serpine1*)^51–54^, and immune reaction (*Il4ra*, *Tnfrsf1a*, *Pgam1*, *Icam1*, *Nfkbia*, *Ifitm3*)^55^. We conclude that esophageal cells growing over an ectopic stroma (eSKIN) remain in an active regenerative state.

Consistent with a transition away from a regenerative state, esophageal cells growing over their own stroma (eEE, **Fig.3E, F** **and** **H**) showed an increased expression of genes associated with epithelial differentiation and keratinization (*Krt13*, *Spink5*, *Sprr1a*, *Klk10*, *Ppl*, *S100a4*, *S100a14*, *Foxq1*)^56–62^, with changes in morphogenic genes known to balance proliferation and differentiation (*Runx1*, *Runx2*)^63^. Other GO terms enriched in eEE-D10 suggested a more matured structural organization of the tissue. These included terms related to cell adhesion (*Dst*, *Dsg2*, *Cldn7*, *Cdh1*, *Gjb4*)^64,65^ cytoskeletal organization (*Tubb3*, *Tubb2a*, *Tuba1a*), and assembly of basement membrane component (*Col4a3*, *Col4a5*, *Col8a1*). The idea of a progressive tissue maturation was further reinforced by the transition from a glycolytic (*Gapdh*, *Gpi1*, *Pfkl*, *Pkm*) to an oxidative signature (*Gpx2*, *Gss*, *Gsr*, *Mgst3*, *Odc1*) from D3 to D10^66,67^. Despite its more differentiated profile, the eEE-D10 showed an increased expression of a subset of developmental genes, such as *Krt8*, *Krt18*, *Krt19*^27,68^ and WNT related genes (*Wnt4*, *Ctnnb1*, *Amer1*, and *Sema3d*), as anticipated for an *in vitro* derived tissue.

The scRNA-seq analysis of the three main transcriptional branches indicated that esophageal cells maintained over an ectopic niche partially retain the regenerative signature found during early re-epithelialization stages (D3).

### 6. Pseudotime analysis reveals the hypoxic signature of esophageal cells transitioning towards skin identity

Having identified the main transcriptional signatures of EE cells exposed to an ectopic niche, we next set out to perform a comprehensive analysis of the molecular changes defining cells undergoing the fate conversion.

To identify transitioning cells, we looked at the expression of typical genes associated with epidermal development or epidermal biology. These markers were enriched in a region of the UMAP represented by the epidermal Host cluster (C18) and the EE-derived cluster 8 (C8) (**Fig.4A** **and** **B**). Cell fate conversion events were therefore restricted to these two clusters, which closely overlap in the UMAP distribution. Interestingly, C8 emerged during early re-epithelialization at D3 and remained present at D10 independent of the stroma used as substrate to establish the esophageal-organ cultures (**SFig.4A**). The signature of this cluster was marked by the expression of genes directly associated with hypoxia, wound healing and cancer (*Pgf*, *Adm2*, *Ero1l*; **SFig.4B**)^34,69–71^. Given that previous *in vivo* scRNA-seq analysis of adult mouse esophagus did not reveal the presence of a C8 equivalent cluster^38^, these results indicate that the C8 population naturally emerges during the normal re-epithelialization process.

**Figure 4.**
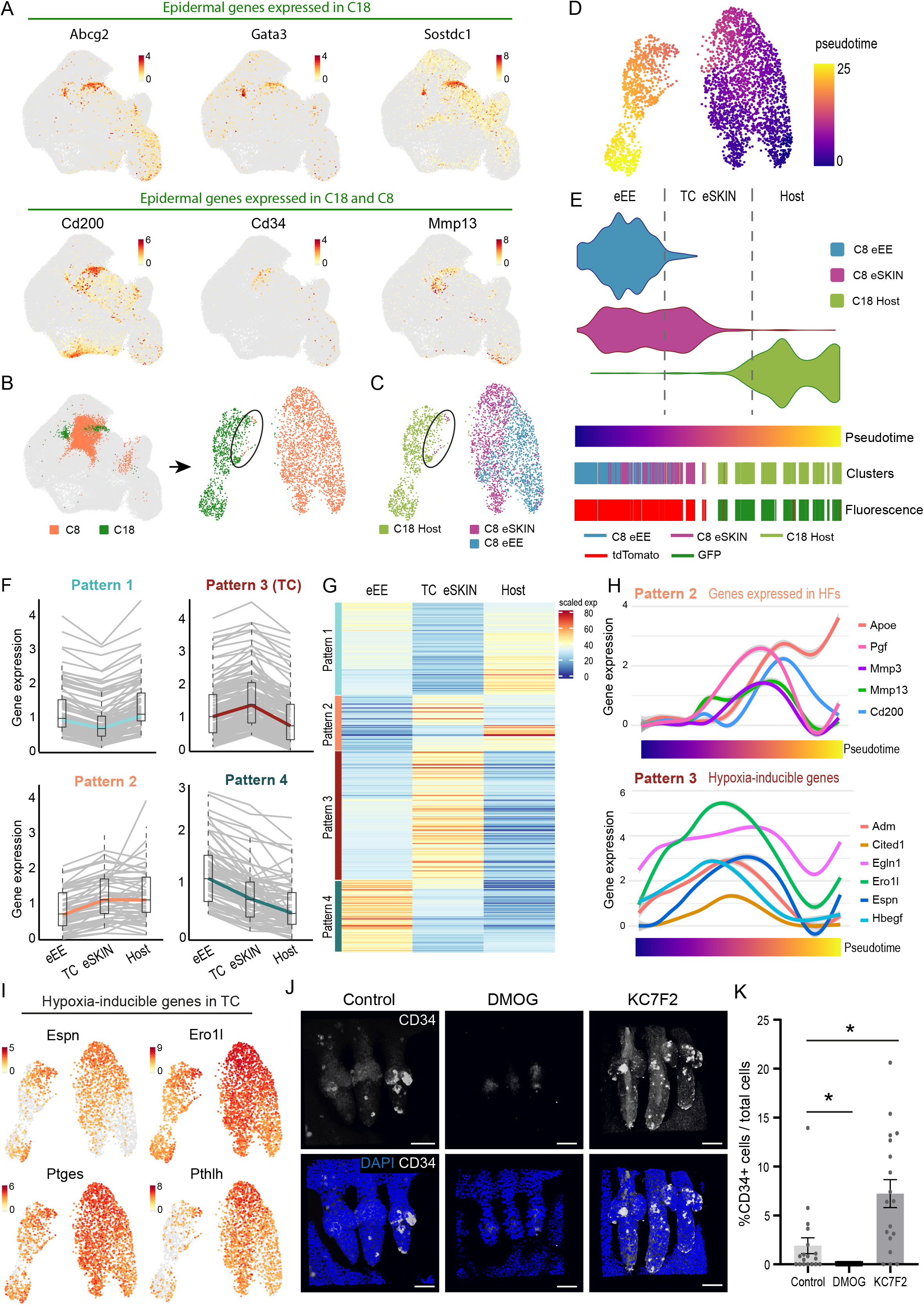
Pseudotime analysis reveals the transcriptional signature of esophageal cells transitioning towards skin identity. **(A)** UMAPs showing expression of classical epidermal genes specifically localized in cluster 18, and hair follicle-related genes localized in clusters 8 and 18. **(B, C)** UMAPs illustrating re-clustering of clusters 8 and 18 at D10. Colors show the identity of the original clusters **(B)** or the sample origin **(C). (D)** Pseudotime analysis of clusters 8 and 18 projected on UMAP space. **(E)** Violin plots representing distribution of cells from different clusters and sample origin along the pseudotime trajectory. Cells were split in three groups along the pseudotime trajectory (dashed lines). Expression-driven rug plots represent the pseudotime trajectory, clusters and predicted tdTomato (esophageal) or EGFP (Host) fluorescent labeling. **(F)** Simplified patterns of gene expression in cells from the three groups defined in **(E)**. Box plots show IQR and whiskers 5^th^-95^th^ percentiles. Colored lines link the median of each group illustrating the pattern dynamics. **(G)** Heatmap representing genes differentially expressed in transitioning cells (TC) *versus* eEE-D10. **(H)** Expression of relevant genes along the pseudotime trajectory from eEE-D10 to Host for genes expressed in hair follicles (pattern 2, top) and hypoxia-induced genes (pattern 3, bottom). **(I)** C8 and C18 UMAPs representing expression of hypoxia-inducible genes up-regulated in transitioning cells (TC). **(J, K)** Images (**J**) and quantification (**K**) of CD34+ cells relative to DAPI+ cells per eHF unit in eSKIN following treatment with DMOG or KC7F2 (activating and inhibiting HIF1α, respectively), as compared to vehicle control (DMSO). Presented as mean ± SEM and analyzed using one-way ANOVA with Kruskal-Wallis multiple comparisons test (*p<0.05 relative to control). n=3 animals. Scale bars 50μm.

Subsequently, we replotted the C8 and C18 separately (**Fig.4B** **and** **C**). In the new UMAP distribution, C8 eEE and C18 Host cells were segregated, with C8 eSKIN cells located between the two (**Fig.4C**). Interestingly, we noticed that cells from eSKIN and Host partially overlapped at the cluster boundary (**Fig.4B** **and** **C**: black ellipse). To explore the trajectory of the identity switch, we performed pseudotime analysis, inferring the differentiation path of esophageal cells (C8) towards skin fate (C18; **Fig.4D** **and** **E**). The pseudotime outlined a trajectory with cells of different origin, i.e. C18 Host and C8 eEE, plotting at opposite ends of the differentiation path, and C8 eSKIN cells transitioning between the two (**Fig.4D** **and** **E**). Next, we examined the gene expression profile along the three main areas defining the pseudotime trajectory, namely eEE, Transitioning cells (TC) and Host (**Fig.4E** dashed lines; see Methods). We identified four major patterns defining changes in gene expression between these areas (**Fig.4F** **and** **G**). Pattern (P) 2 contained genes with sustained increased expression in host cells. As anticipated, this included genes typically expressed in HFs (*Cd200*, *Apoe*, *Mmp9*, *Sparc*)^72–75^, as well as other genes involved in extracellular matrix (ECM) remodeling, migration and invasion (*Fhl2*, *Cst3*, *Aqp3*, *Cav1*, *Cd63*, *Mmp3*, *Mmp13*) (**Fig.4H**; **SFig.4C**). P4 reinforced the notion of eSKIN cells changing cell fate identity, by showing a reduction in epithelial genes expressed at low levels in Host cells (*Krt7*, *Krt13*, *Pitx1*, *S100a4*, *Wfdc18*; **Fig.4F** **and** **G**; **SFig.4C and E**)^58,76^. P3 represented the most biologically relevant pattern, containing genes specifically upregulated in TCs undergoing cell fate conversion (**Fig.4F** **to** **H**; **SFig4C**). GO enrichment analysis for these genes revealed processes previously shown to be associated with cell fate plasticity^77,78^. This included terms defining an active tissue remodeling signature i.e. wound healing, invasion, EMT and tumor progression (*Hbegf*, *Serpine1*, *Bsg*, *Plod2*, *Pthlh*, *Cited1*)^79–85^. Other genes denoted the active regenerative state of TCs, including those associated with ECM reorganization (*Lamc2*, *Flrt3*, *Adam8*) and glycolytic metabolism (*Aldoc*, *Gapdh*, *Gpi1*, *Pgam1*, *Pkm*, *Tpi1*) (**SFig.4C and D**). Interestingly, the hypoxic profile, already defining cells in C8, was notoriously more accentuated in cells transitioning towards skin identity (TC), expressing particularly high levels of genes regulated by HIF1a (*Adm*, *Bnip3*, *Ddit4*, *Egln1*, *Egln3*, *Ero1l*, *Pgk1*, *Ptges*, *Vegfa*; **Fig.4H and I**)^69,86–88^. Consistent with TCs being at a more regenerative and less differentiated state compared to cells of established identity (eEE and Host), genes associated with protein biogenesis (*Ddx21*, *Gnl3*, *Ncl*, *Nop56*, *Nop58*, *Rexo2*)^89^ and oxidative metabolism (*Mgst1*, *Mgst2*, *Prdx6*)^76^ were downregulated at the transition^66,90^ (Pattern 1; **Fig.4F** **and SFig4C**).

Our scRNA-seq analysis indicates that the transcriptional signature of esophageal cells transitioning towards skin identity show a regenerative profile activated during early re-epithelization stages, independent on the origin of the niche. Interestingly, the pseudotime analysis highlights a clear transcriptional shift, specific of transitioning cells, towards a strong hypoxic signature marked by the expression of HIF1a downstream targets.

### 7. HIF1a inhibition promotes esophageal-to-skin identity switch

Our scRNA-seq analysis revealed that esophageal cells transitioning towards skin identity showed a transient regenerative signature, marked by the expression of hypoxia associated genes and, in particular, HIF1a downstream targets. To confirm the relevance of this pathway in cell fate conversion, we treated our heterotypic cultures with a known HIF1a activator (DMOG) or inhibitor (KC7F2)^91,92^, and measured CD34 staining in the newly formed eHFs as a read out of identity transition. The results showed a significant increase in the % of CD34+ cells within eHFs upon inhibition of HIF1a, with the opposite effect in response to the activation of the pathway (**Fig. 4J** **and** **K**). A similar pattern was found in the percentage of eHFs positively stained with CD34 (26% in control; 61.7% in KC7F2; 12.9% in DMOG). Consistent with the transient hypoxic signature observed along the conversion trajectory (**Fig.4H** **and** **I**), our functional assay suggests that down-regulation of HIF1a signaling pathway promotes the exit from an identity transition state and favors fate re-specification as instructed by the ectopic stroma.

## Discussion

For many decades, squamous epithelial cells were thought to be structured in a rigid hierarchy where stem cells were responsible for tissue maintenance and regeneration, while committed and differentiated cells only contributed to the formation of the epithelial barrier^5–7^. Mounting evidence over the last two decades has now established that epithelial cell behavior present a remarkable plasticity^8^. Following injury, differentiating or lineage committed cells have the ability to reacquire stem cell-like behavior increasing the regenerative response of the tissue^12,14,17,93^.

Over the years, efforts to understand the rules governing cell fate choices have unveiled the key role of the niche in modulating epithelial behavior. Early tissue recombination studies in chicks uncovered that epidermal cells, independently of their region of origin, are able to form feathers or scales if instructed to do so by dermis bearing those structures^24,94^. More recent work in the mouse skin has shown that changes in the mesenchymal-epithelial cross-talk during developmental stages can direct epidermal cells to re-specify their appendage forming capacity, switching between HF and sweat gland identity^25^. Further, in an effort to explore whether cell fate conversion is a widespread phenomenon beyond the epidermis, a set of epithelial tissues of non-skin origin have been shown to have the potential to switch towards HF identity when exposed to the developing dermis of hair-bearing skin^21^. Yet, the mechanisms governing plasticity, particularly in adult tissues, remain unclear.

Here we developed a 3D heterotypic culture system to investigate the mechanisms regulating epithelial plasticity in adult tissues. Using this approach, we have explored the temporal conversion of esophageal cells into hair follicles when exposed to the foreign stroma of the adult skin. Single-cell transcriptional profiling at different time points throughout the re-specification process provided insight into the mechanisms modulating adult cell fate plasticity. In particular, our results showed that the regenerative signature, acquired during early re-epithelialization stages, is retained by those cells undergoing the identity switch at later time points. Cell fate trajectory analysis further revealed that the regenerative signature of transitioning cells is marked by a strong hypoxic component, expressing high levels of HIF1a downstream targets. Interestingly, recent work has shown that the hypoxic niche of the upper hair follicle promotes the stem cell state^95^, raising the question as to whether this environment may be playing a role during the esophageal cell fate conversion. Functional validation experiments demonstrated that HIF1a inhibition forces cells away from the transitioning state, in which most cells persist, favoring esophageal cell fate conversion as instructed by the ectopic microenvironment. Our results unveil the central role of HIF1a pathway in modulating adult epithelial cell fate plasticity. Future studies will be required to investigate whether the molecular changes identified using our *ex vivo* system are important for other clinically relevant models, such as wound healing and cancer, where plasticity is known to operate^8,9,11,13,93^.

## Methods

### Mouse strains and induction of allele

All mouse experiments were approved by the local ethical review committees at the University of Cambridge, and conducted according to Home Office project license PPL70/8866 at the Wellcome Trust-Medical Research Council Cambridge Stem Cell Institute, Cambridge University.

To identify the cell origin (epithelial or stromal) in 3D heterotypic cultures the following fluorescent reporter mouse lines were used: mTmG (R26mTmG; stock #007676, Jackson Laboratory^96^) and nTnG (R26nTnG; stock #023537, Jackson Laboratory^39^), which constitutively express tdTomato localized in the cell membrane or the nuclei, respectively; H2B-EGFP (CAG::H2B-EGFP; stock #006069, Jackson Laboratory^97^) with constitutive GFP nuclear expression; and Lgr5-EGFP (Lgr5-EGFP-IRES-creERT2; stock #008875, Jackson Laboratory^98^) with an EGFP cassette targeted to the 3’ untranslated region of the Lgr5 gene.

To visualize the cell proliferative state of cells, the Fucci2a mouse line (R26Fucci2a, kindly provided by Ian J. Jackson (Mort et al., 2014)), which incorporates constitutive expression of cell cycle reporter proteins (G1 marked by mCherry-hCdt1, and S/G2/M marked by mVenus-hGem), was used.

For lineage tracing analysis, R26-CreERT2 animals (stock #008463, Jackson laboratory (Ventura A, 2007)) were crossed onto R26R-Confetti (stock #017492, Jackson laboratory (Clevers H, 2010)) to generate R26-CreERT;R26R-Confetti mice, which following Cre-mediated recombination express one of four fluorescent reporter proteins (YFP, GRP, RFP or CFP) in scattered cells. R26-CreERT;R26R-Confetti mice were induced by a single intraperitoneal (i.p.) injection of tamoxifen (3 mg per 20 g of body weight).

C57BL/6J mice (strain code, 632) and nude athymic mice (strain code, 490) were purchased from Charles River, UK. All experiments comprised mixed male and female animals, apart from RNA sequencing experiments for which only male animals were used in order to avoid cofounding effects due to estrous cycle. All animal cohorts were adults between 8-25 weeks of age.

### Histology

Hematoxylin and Eosin (H&E) staining was performed in 10μm paraffin-embedded sections by the Histology Core Service at Cambridge Stem Cell Institute and imaged using Apotome Imaging System (Zeiss).

### Heterotypic 3D cultures

In order to study cell fate changes in esophageal epithelium (EE) when exposed over the skin stroma, we adapted a previously described ex vivo explant technique^19^. As a control, esophageal epithelium was grown over its native stroma (eEE). Unless otherwise stated esophageal tissues were tdTomato+ (mTmG or nTnG mouse lines), and tissues of skin origin are EGFP+ (from H2B-EGFP line). Skin and esophageal stroma – dermis and submucosa, respectively – were dissected away from the epithelial tissue layer after 3-4 hours incubation in 5mM EDTA at 37°C. The stroma was cut in 9 × 7 mm pieces for the dermis and 5 × 10 mm pieces for the submucosa and placed onto transparent ThinCertTM inserts of 0.4 μm pore size (Greiner Bio-One Ltd; Cat#657641). Immediately afterwards, two strips of esophageal epithelia (5 × 1 mm) were laid on the stroma. The mounted explants were allowed to settle for 5 minutes at 37°C to ensure attachment, and then covered with minimal medium containing one part DMEM (4.5 g/L D-Glucose, Pyruvate, L-Glutamine), one part DMEM/F12, supplemented with 5% fetal calf serum, 5 μg/ml insulin, 0.18 mM adenine, 5-10 μg/ml transferrin and 5% Penicillin-Streptomycin. Cultures were grown for up to 10 days at 37°C and 5% CO2, replacing the medium on alternate days. During this period of time the stroma was re-epithelialized by esophageal cells. The newly formed epithelial sheet was either cryosectioned or wholemounted by peeling away from the stroma following 2 hours incubation in 5mM EDTA at 37 °C. Newly formed epithelium of esophageal origin grown over denuded dermis (eSKIN; eHF, Hair Follicle; eIFE, Interfollicular Epidermis). Newly formed epithelium of esophageal origin grown over denuded submucosa (eEE)

As an in vitro skin control, the dermis was cultured in the absence of esophageal tissue explants. This system allowed epidermal re-growth by sporadic epidermal cells remaining in the dermis. The resulting epithelial layer was wholemounted as described (these control samples include sSKIN, sHF, sIFE).

### Tissue wholemount preparation and immunofluorescence

Wholemounts from the middle third EE and tail skin were obtained as follows: Tissues were cut into pieces of approximately 5 × 8 mm and incubated for 3-4 hours in 5mM EDTA at room temperature. The epithelium was then gently peeled away from the underlying tissue and fixed in 4% paraformaldehyde (PFA) in PBS for 30 minutes. For staining, wholemounts were blocked in staining buffer (0.5% Bovine Serum Albumin, 0.25% Fish skin gelatin, and 0.5% Triton X-100 in PBS) with 10% donkey serum for 1 hour at 37 °C. Wholemounts were then stained with primary antibodies (see below) in staining buffer overnight at 4 °C followed by washing for 2 hours with 0.2% Tween-20 in PBS. Samples were incubated with secondary antibodies in staining buffer for 3 hours at room temperature before washing as above. Cell nuclei were counterstained with 1 μg/ml DAPI in PBS and samples mounted in 1.52 RapiClear mounting media for imaging.

For staining of tissue cryosections, fixed samples were embedded in optimal cutting temperature compound (OCT), cut at 150 μm and placed onto poly-L-lysine coated glass slides. Cryosections were stained and mounted following the wholemount protocol.

The following primary antibodies were used: CD34 (Rat, 1:100, BD Bioscience), CD45 (Rat, 1:200, BioLegened), CD49f (integrin α6, Rat, 1:200, BioLegend), KRT4 (Mouse, 1:2000, Vector Laboratories), KRT14 (Rabbit, 1:1000, BioLegend), KRT17 (Rabbit, 1:250, Cell Signalling), KRT24 (Rabbit, 1:500, Atlas) and Sox2 (Rat 1:200, eBioscience).

### Confocal imaging and analysis

Images were acquired using an inverted Leica SP5 confocal microscope (Leica Microsystems) with standard laser configuration. Typical settings for z-stack image acquisition included optimal pinhole, line average 3, bi-directional scan with speed of 400 Hz and a resolution of 1024 × 1024 pixels using 40x objective. Images were typically acquired with digital zoom 3x for EE and IFE epithelium and 1x for HFs. Confocal images were reconstructed and analysed using Volocity 5.3.3 software (PerkinElmer).

For cell quantification analysis a minimum of 18 HF units were imaged and quantified (n=3 animals) and representative images are shown. To calculate HF markers expression rate, a minimum of 150 eHF units (n=3 animals) were examined on live scanning mode and representative images taken.

### eHF-formation efficiency

9 × 7 mm tail skin pieces from Lgr5-EGFP-IRES-creERT2 reporter mice were incubated in 4 μg/ml DAPI in PBS, followed by clearing in 1.52 RapiClear and tile-scanned on a confocal microscope. Immediately after imaging, samples were washed in PBS and incubated in 5mM EDTA for 4 hours at 37°C. The epidermis was dissected away from the dermis and esophageal tissue from mTmG mice laid on top. Heterotypic cultures were grown for 10 days and full thickness tissue fixed in 4% PFA for 1 hour. Wholemounts were then washed three times in PBS and mounted in 1.52 RapiClear mounting media for imaging. GFP+ and mTomato+ HF structures – before and after culture, respectively - were quantified using xyz views of rendered confocal tiled images. eHF formation efficiency in vitro was calculated as the ratio of eHF to the number of original HFs in the in vivo skin piece. 3 samples were quantified (n=3 animals).

### In vivo transplantation

Non-cultured heterotypic grafts were prepared with dermis obtained from H2B-EGFP animals and esophageal tissue from mTmG or nTnG mice, unless otherwise stated. To reduce tissue dehydration during the procedure, one single esophageal explant was laid on the dermis and kept in a humidified environment until transplantation. Female nude athymic mice were used as recipients. In a typical experiment, anesthetized recipients underwent a single excision on the dorsal shoulder using sterile 8 mm diameter biopsy punch. Immediately afterwards, a heterotypic graft (8 mm diameter) was fitted and adhered to the surrounding tissue with Vetbond tissue adhesive (3M). The area was further sutured using steri-strip wound closure strips (3M) and covered with Tegaderm dressing film (3M) to protect the tissue implant. Animals were monitored thereafter, and skin tissue harvested up to 4 weeks post-transplant.

### Hair reconstitution assay

The experiments were performed similarly as previously described^34,36^. Back skin from postnatal day 2 (P2) H2B-EGFP mice was dissected and floated epidermal side up in 10ml 0.25% trypsin overnight at 4°C. After washing with PBS, the epithelium was peeled off and the dermis was minced and incubated in 0.25% collagenase for 25min at 37°C. The dermal cell suspension was then filtered with a 100 micron strainer and the HF buds were pelleted by centrifugation for 5min at × 100g. The supernatant was transferred to a new tube and centrifuged for 5min at × 300 g. Dermal cells were resuspended in PBS, and kept on ice until transplantation. To isolate esophageal epithelial cells, esophagus from adult nTnG mice were cut in 4 pieces and incubated in 0.5mg/ml Dispase for 15min at 37°C. The EE was peeled, minced with a scalpel and transferred to a new tube with fresh Dispase. After 5min of incubation at RT, EDTA was added at a final concentration of 5mM to inhibit Dispase activity. Cells were mixed by pipetting and filtered throught 30μm strainers. EE cell suspensions were centrifuged 5min at x 300g, resuspended in PBS and kept on ice until transplantation. Isolated esophageal epithelial cells were mixed with neonatal dermal fibroblasts and surgically implanted on the dorsal fascia of recipient nude mice after a full-thickness 8mm diameter biopsy punch. Reconstituted hair follicle appendages were observed and harvested for immunofluorescence analysis 6 weeks post grafting. Typical number of transplanted cells was 2×106 EE cells and 4×106 neonatal skin cells.

### Confetti labelling

Tamoxifen (Sigma) was dissolved in sunflower seed oil at a concentration of 30 mg/ml. Ubiquitous low frequency recombination of the confetti cassette was achieved by inducing animals with an intraperitoneal dose of 3mg per 20g of mouse body weight. 48-hours following induction, animals were harvested and the esophagus was used for cultures and to assess induction efficiency.

### Single-cell RNA isolation and library preparation

Heterotypic 3D cultures were prepared using EE from nTnG mice and esophageal stroma or skin dermis from H2B-EGFP mice. eEE and eSKIN were harvested after 3 and 10 days and the epithelia were carefully peeled following incubation with 50mM EDTA for 15min at 37⍰C. Peeled epithelia were washed with PBS and incubated with 0.5mg/ml Dispase (Sigma) for 5min. EDTA was then added to the samples at a final concentration of 5 mM, and the suspension diluted 1/5 by adding FACS buffer (FB; 2% heat-inactivated Fetal bovine serum (Life Technologies; 26140079), 25 mM HEPES (Life Technologies; 15630056)) in order to reduce Dispase activity. A single-cell suspension was obtained by filtering through a 30 μm cell strainer. Cells were then centrifuged at 300 g for 10 minutes at 4°C, and resuspended in FB containing 1U/μl RNAse Inhibitor. Samples were sorted on a BD FACSAria™ III cell sorter. Cell suspensions were incubated with DAPI for 1 min prior sorting and first gated against DAPI to exclude dead cells, and then with forward and side scatters to gate for singlets. Tissue-origin gating was performed with tdTomato and EGFP. scRNA-seq libraries were generated using 10X Genomics kits (Single Cell 3’ v3) at the CRUK-CI Genomics Core Facility of the CRUK Cambridge Institute, using 9,000 cells per sample.

3 biological replicates for eEE-D3, eSKIN-D3, and eEE-D10, and 5 biological replicates for eSKIN-D10 summed up to 14 10x libraries. Each biological replicate consisted of pooled cultures from 4-5 wells. The libraries were multiplexed and sequenced on 4 lanes Illumina NovaSeq6000 S2 flow cells.

### Single-cell RNA-seq analysis

Pre-processing quality checks were performed on the R2 reads using FastQC (https://www.bioinformatics.babraham.ac.uk/projects/fastqc). The data was then processed using the cellranger software (v3.1.0) using standard alignment (using the genome assembly GRCm38.97 of M musculus) and default filtering parameters.

Upper and lower bounds on the distributions of counts, features, mitochondrial and ribosomal RNA were used to remove outlier cells (cells were accepted if their sequencing depth was over 8750, the number of expressed genes was between 2000 and 8000, and the percentage of mitochondrial and ribosomal DNA was lower than 15% and 45%, respectively). Mitochondrial and ribosomal genes were subsequently removed from the matrix. The raw expression matrix was normalized using sctransform (http://dx.doi.org/10.1186/s13059-019-1874-1); the downstream was performed using the Seurat R package (v3.2.2)^99^.

Dimensionality reductions (PCA and UMAP), as well as clustering were also performed using the Seurat R package, with the number of clusters being the optimal selected by the software with default parameters. Cells expressing fibroblast or immune marker genes including Col1a2, Pdgfra or Ptprc were considered as non-epithelial contaminants and filtered out of the analysis. For the subsequent analysis, we focused on the 1000 most abundant genes. Markers for each cluster were identified by testing for differential gene expression (in Seurat). The above steps were performed both for all the cells in the data, but also individually for each sample set (eEE-D3, eEE-D10, eSKIN-D3, eSKIN-D10).

Clustering stability was evaluated using a PAC analysis (https://doi.org/10.1038/srep06207). Enrichment analysis on the markers identified per cluster was performed using the g:profiler (https://biit.cs.ut.ee/gprofiler/gost) R package (v0.1.8)). Cells marked with the fluorescent markers EGFP and tdTomato were identified through sequencing. To avoid high similarity regions between EGFP and tdTomato, we selected, based on a ClustalOmega alignment, 3 and 5 sub-transcripts respectively, with less than 20% identity between the two markers. Using STAR aligner (v2.5.2a), with relaxed mapped length parameters, we aligned the reads across all samples to the selected sub-transcripts; the expression levels across the sub-transcripts was defined as the algebraic sum of incident reads. Next we selected ~70 cells with the highest, independent, expression of red or green markers, respectively; these two sets were used to train a classifier, part of a semi-supervised model that extends the transcriptomic signatures of the EGFP and tdTomato cells to all unlabeled cells i.e. cells with no or little quantification of the fluorescent markers. The discriminative genes were selected using a random forest model (the top 50 genes with the highest variation of gini index) and standard statistical tests i.e. t-test, Kolmogorov-Smirnov and Mann–Whitney U; genes with adjusted p-value less than 0.05 were selected as discriminative. The final set of markers, used subsequently for inferring labels, was obtained by intersecting the two approaches. Next, the identity of the unlabeled cells was predicted. For each unlabeled cell, we calculated a correlation-based distance (Pearson and Spearman approaches yielded comparable results), on the selected markers, to all tdTomato and EGFP cells used for the original classifier. The two distributions of distances were compared using a minimum IQR approach i.e. if the distance between the red and green distributions on the proximal quartiles was larger than the average standard deviations, then the cell was labelled according to the distribution closer to 0, otherwise the cell was labelled “unknown”.

After identifying cluster 18 as Host EGFP expressing cells, the remaining clusters were manually annotated based on gene expression of known cell cycle, epithelial basal and differentiated marker genes (Fig.S3H and I). We then focused on differences in gene expression of the of basal cells in the three major transcriptional profile branches (D3, eSKIN-D10 or eEE-D10). To this end, clusters of annotated basal cells were assigned to one branch if >75% cells belonged to the correspondent sample origin. Clusters 3 and 8 had an equivalent contribution of the three sample origins and therefore cells in these clusters were split in the three branches depending on their origin. Next, we identified differentially expressed genes (DEGs) for each of the comparisons between branches. The total number of DEGs was 567. We then scaled the expression value for each gene in the [0,100] range using an affine transformation and summarised the scaled expression levels in a clustered heatmap. Of the resulting 6 patterns, we selected those showing a similar transcriptional signature between D3 and eSKIN-D10, and performed an enrichment analysis. using g:profiler (https://doi.org/10.1093/nar/gkz369), against the standard GO terms, and the KEGG (https://doi.org/10.1093/nar/28.1.27) and Reactome (https://doi.org/10.1093/database/baz123) pathway databases. The observed set comprised of the selected genes, the background set comprised of all genes expressed in the dataset.

We then focused on identifying cell fate conversion events within cluster 8, enriched in epidermal marker genes, and cluster 18 (Host). To that end, we re-clustered cells from C18, and from eEE-D10 and eSKIN-D10 in C18, time point were the cell identity switch had been described experimentally. In the new UMAP space, eSKIN-D10 from C8 were found between eEE-D10-C8 and C18. Thus, we performed a pseudotime analysis using the monocle3 R package (v0.2.3.0) to project the expression profile of relevant genes along the cell fate conversion journey. As expected from the UMAP visualization, the transitioning cells (TCs) were located in the middle of the pseudotime trajectory, between the eEE and the Host cells. To examine in more detail the changes occurring in these TC, we defined three equidistant groups along the pseudotime axis, summarized the expression of cells belonging to these groups and identified the corresponding DEGs by finding the markers of each pairwise comparison between groups. We obtained 1173 DEGs. To focus on the exclusive features of the TC, we selected the genes that had >0.2log2FC between the eEE-D10 and the TC, and presented them in a clustered heatmap using the Pearson correlation for the clustering to reveal patterns of expression. The clustered dendrogram was cut to identify 4 patterns. We run enrichment analysis using g:profiler (https://doi.org/10.1093/nar/gkz369), against the standard GO terms, and the KEGG (https://doi.org/10.1093/nar/28.1.27) and Reactome (https://doi.org/10.1093/database/baz123) pathway databases. The observed set comprised of the selected genes, the background set comprised of all genes expressed in clusters 8 and 18. Terms and genes were manually curated the lists, in accordance to the biological context of the project.

### Hypoxia treatments

Heterotypic cultures were treated with the HIF1a translation inhibitor KC7F2 (10 μM; SML1043, Merck) or the HIF1a protein stabilizer DMOG (1mM;71210, Cayman) during a period of 8 days. Control cultures were treated with DMSO vehicle alone. Cultures were grown in a normoxic humidified incubator at 37°C, 5% CO2 and medium changed on alternate days.

### Statistical Analysis

Number of animals used for each experiment are indicated in the figure legends. Experimental data are expressed as mean values ± SEM unless otherwise stated. Differences between groups were assessed by using two-tailed unpaired t-test, one-way or two-way analysis of variance (ANOVA) as indicated in figure legends. ANOVA based analysis was followed by Tukey’s test for multiple comparisons. Statistical differences between groups were assessed using GraphPad Prism software. No statistical method was used to predetermine sample size. Experiments were performed without methods of randomization or blinding.

## Supporting information

Supplementary file Bejar MT, Jimenez-Gomez P et al.

## Author Contributions

M.T.B, P.J.G, F.J.C.N and M.P.A designed, validated and conducted experiments; I.M (Illias) performed single-cell RNA sequencing data analysis supervised by I.M (Irina). S.H, F.J.C.N, and B.G guided the experimental design for single-cell RNA and provided advice on data analysis. B.C and B.D.S supervised parts of the study and provided expertise in the epithelial stem cell field. M.T.B, P.J.G, B.C and M.P.A conceived the project, supervised experiments, and wrote the manuscript with input from authors. Funding acquisition by M.T.B, B.G, B.D.S, and M.P.A. M.P.A supervised the study.

## Acknowledgments

We thank members of the Alcolea’s lab for comments and suggestions; Oriana Oniciuc at the bioinformatics facility at JCBC for her contribution to the scRNA-seq analysis; the staff of the University Biomedical Services Gurdon Institute; Peter Humphreys in imaging core facilities at JCBC; Alex Sossick, Richard Butler and Nicola Lawrence at the Gurdon Institute Imaging Facility; Ian J. Jackson (Fucci2a); NIHR Cambridge BRC Cell Phenotyping Hub for FACS support; Cambridge Cancer Institute Genomics Facility for sequencing; the Jeffrey Cheah Biomedical Centre (JCBC) core facilities. This work was mainly supported by funding from the Wellcome Trust and The Royal Society (105942/Z/14/Z to M.P.A), and a core support grant from the Wellcome and MRC to the Wellcome-Medical Research Council Cambridge Stem Cell Institute. M.T.B received funding from the European Union’s Horizon 2020 research and innovation programme under the Marie Sklodowska-Curie grant agreement No 794664 (OESOPHAGEAL FATE). P.J.G was supported by a Wellcome PhD stutentship (102160/B/13/Z) The work also received support from Isaac Newton Trust (Research Grant 16.24(e) to M.P.A); Human Frontier Science Program (LT000092/2016-L) and Basic Science ResearchProgram (NRF-2014R1A6A3A01005675) to S.H; Wellcome Trust (206328/Z/17/Z to F.J.C.N, and B.G); B.D.S. acknowledges funding from the Royal Society E.P. Abraham Research Professorship (RP\R1\180165), the Wellcome Trust (219478/Z/19/Z) and MRC (MR/V005405/1).

