## Supplementary file Bejar MT, Jimenez-Gomez P et al. for "Defining the transcriptional signature of esophageal-to-skin lineage conversion"

### Figure S1

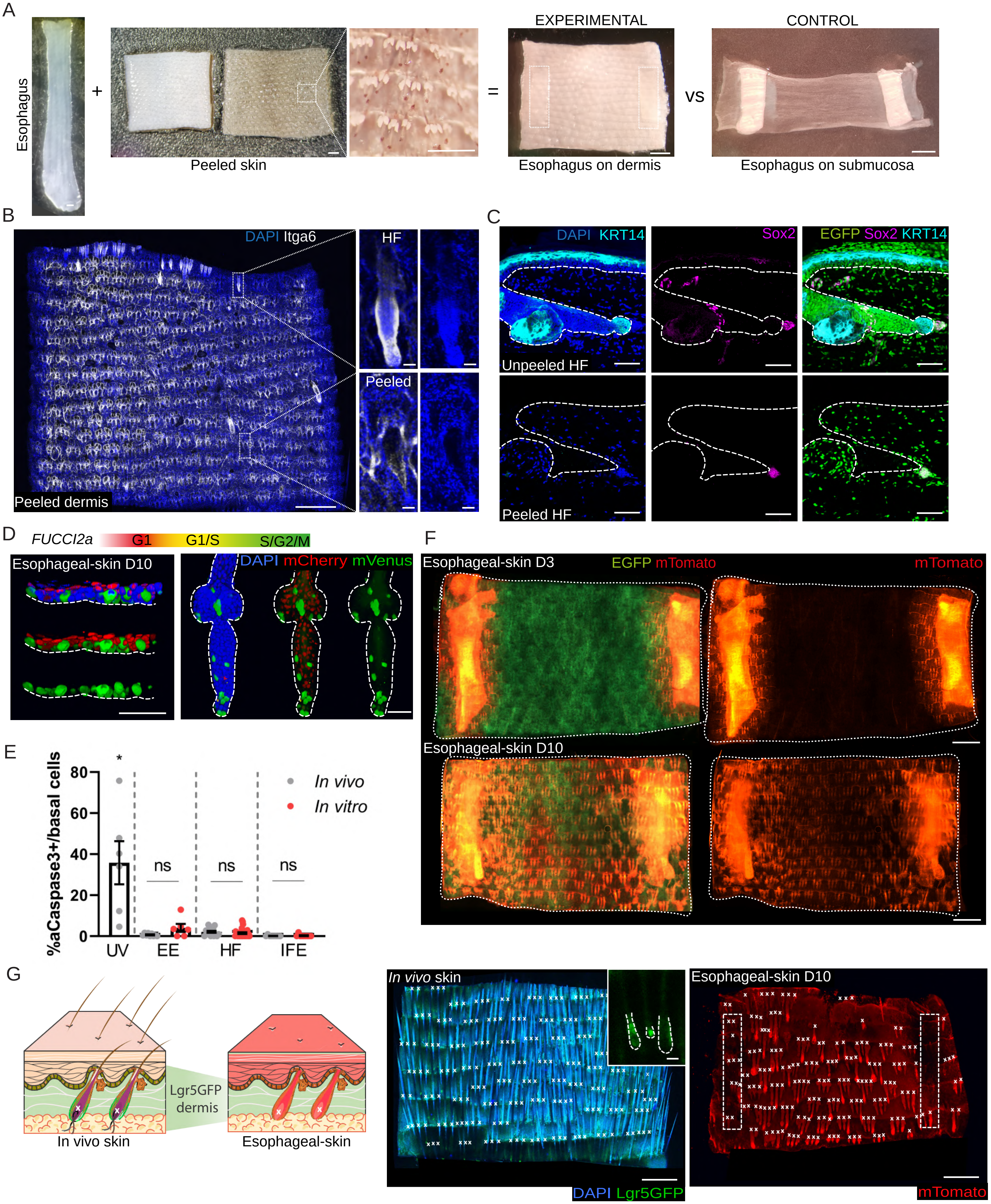
A
Esophagus
+
Peeled skin
=
EXPERIMENTAL
Esophagus on dermis
vs
CONTROL
Esophagus on submucosa
B
DAPI Itga6
Peeled dermis
HF
Peeled
C
DAPI KRT14
Sox2
EGFP Sox2 KRT14
Unpeeled HF
Peeled HF
D
FUCCI2a
G1
G1/S
S/G2/M
Esophageal-skin D10
DAPI mCherry mVenus
E
%aCaspase3+/basal cells
80
60
40
20
0
*
ns
ns
ns
UV
EE
HF
IFE
In vivo
In vitro
F
Esophageal-skin D3
EGFP mTomato
mTomato
Esophageal-skin D10
G
Lgr5GFP dermis
In vivo skin
Esophageal-skin
In vivo skin
DAPI Lgr5GFP
Esophageal-skin D10
mTomato

**Supplementary Figure 1. 3D heterotypic culture set up and characterization. Related to Figure 1. (A)** 3D heterotypic culture composite set up. Esophageal and skin epithelia are separated from stroma (dermis for skin and submucosa for esophagus). Inset shows peeled skin epidermis containing hair follicles. Esophageal epithelial strips (white dashed lines) are laid on isolated skin dermis or esophageal submucosa (as controls) and cultured for up to 10 days. See also **Fig.1B**. **(B)** 3D rendered confocal images showing peeled dermis from a wild-type mouse stained with DAPI (blue) and integrin α6 (itga6, greyscale). Insets show a representative empty hair follicle (HF) socket (bottom images) and sporadic HFs remaining in the dermis (top images). **(C)** Confocal images of skin cryosections from green H2B-EGFP mice before (top) and after (bottom) peeling the epidermis. The epithelial layers (positively stained with KRT14, white) are removed in the peeling process, whereas dermal papillae (stained with Sox2, magenta) remain in the HF socket. **(D)** Schematic at the top illustrates the expression pattern of fluorescent cell cycle reporter proteins in the Fucci2a mouse model (mCherry, G1 cells; mVenus, S/G2/M cells). Esophagi from Fucci2a mice were grown for 10d in 3D heterotypic cultures. 3D rendered confocal images show side view of esophageal-derived interfollicular epidermis (left panel) and esophageal-derived HF (right panel). n=3 animals. **(E)** Quantification of activated-Caspase3+ cells in esophageal-derived skin *in vitro* and in tissue wholemounts *in vivo* relative to total basal cells. Ultraviolet (UV) irradiation of EE was used as a positive staining control. Presented as mean ± SEM and analyzed using one-way ANOVA with Kruskal-Wallis multiple comparisons test (*$p<0.05$ relative to *in vivo* samples, ns = not significant). n=3 animals. **(F)** Representative images of wholemounts from 3D heterotypic cultures at day 3 (D3, top) and day 10 (D10, bottom) as in **Fig.1B**. Donor green H2B-EGFP mouse dermis, and esophagus from mTmG (tdTomato, red) mouse line. **(G)** Left: schematic exemplifying the method to quantify the re-epithelization efficiency of the heterotypic culture. Donor green Lgr5-EGFP mouse dermis, and esophageal epithelium from mTmG (tdTomato, red) mouse line. Right: 3D images of *in vivo* skin wholemount (left image, inset shows *in vivo* Lgr-5GFP HFs, delineated by white dashed lines) and esophageal-derived skin from heterotypic culture at D10 (right image, white dashed lines delineate esophageal strips). The number of individual HF units (white crosses) were measured before (green) and after (red) peeling to measure the efficiency of esophageal-derived HF formation. n=3 animals. **Scale bars.** S1A-B, S1F-G (1 mm), insets (50 µm); S1C-D (50 µm).

Figure S2

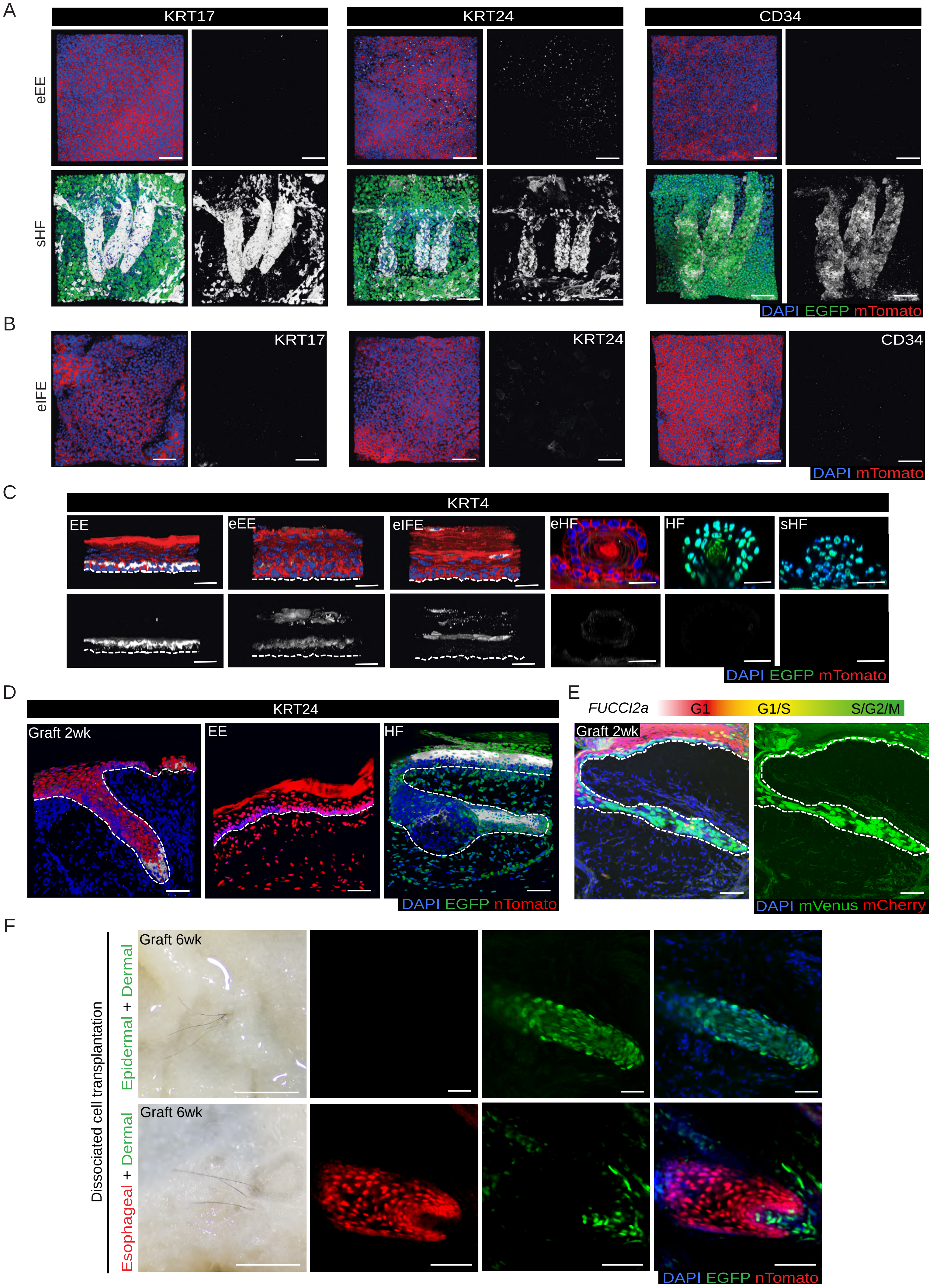
A
KRT17
KRT24
CD34
eEE
sHF
DAPI EGFP mTomato
B
eIFE
KRT17
KRT24
CD34
DAPI mTomato
C
KRT4
EE
eEE
eIFE
eHF
HF
sHF
DAPI EGFP mTomato
D
KRT24
Graft 2wk
EE
HF
DAPI EGFP nTomato
E
FUCCI2a
G1
G1/S
S/G2/M
Graft 2wk
DAPI mVenus mCherry
F
Dissociated cell transplantation
Epidermal + Dermal
Esophageal + Dermal
Graft 6wk
Graft 6wk
DAPI EGFP nTomato

**Supplementary Figure 2. *In vitro* and *in vivo* validation of esophageal-to-skin cell fate conversion. Related to Figure 2. (A)** Representative 3D rendered confocal z-stacks showing *in vitro* esophageal epithelium (eEE) and *in vitro* skin-derived HF (sHF) as *in vitro* controls for **Fig.2A**. n=3 animals. **(B)** Representative confocal z-stacks showing basal view of esophageal-derived IFE (eIFE). n=3 animals. Labeling in **(A,B)** show HF markers KRT17, KRT24 and CD34 in greyscale, as indicated. **(C)** Confocal images showing single plane images for HFs and side views for other tissues. Esophageal-derived skin (eIFE, eHF); *in vitro* (eEE, sHF) and *in vivo* controls (EE, HF) n=3 animals. Labeling shows esophageal differentiation marker KRT4 in grey scale. **(D)** Cryosections of heterotypic *in vivo* transplantation 2-week graft (left), compared to esophagus (middle) and HF (right) controls. **(E)** Schematic illustrates expression pattern of fluorescent reporter proteins in Fucci2a mouse model. Esophagi from Fucci2a mice were used for heterotypic transplantation *in vivo.* Recipient animals were collected 2 weeks post-transplant. Blue, DAPI; Red, mCherry (G1); Green, mVenus (S/G2/M). n=2 animals. **(F)** Images show newly-formed hairs in nude recipient mice following a 7 week cell grafting (mix of dissociated cells from neonate skin (EGFP) and adult EE cells (tdTomato; nTnG) subcutaneously transplanted). Confocal images show HFs derived from epidermal cells (top panel) as well as esophageal cells (bottom panel). **Scale bars.** S2A-S2E (50 µm); S2F (hair images, 1 cm; confocal images 50 µm). **Fluorescent reporters and stainings.** Unless otherwise indicated tissues of esophageal origin in red (mTmG **(A-C)** and nTnG **(D-F)** mouse line), and skin tissues in green (H2B-GFP mouse line).

#### Figure S3

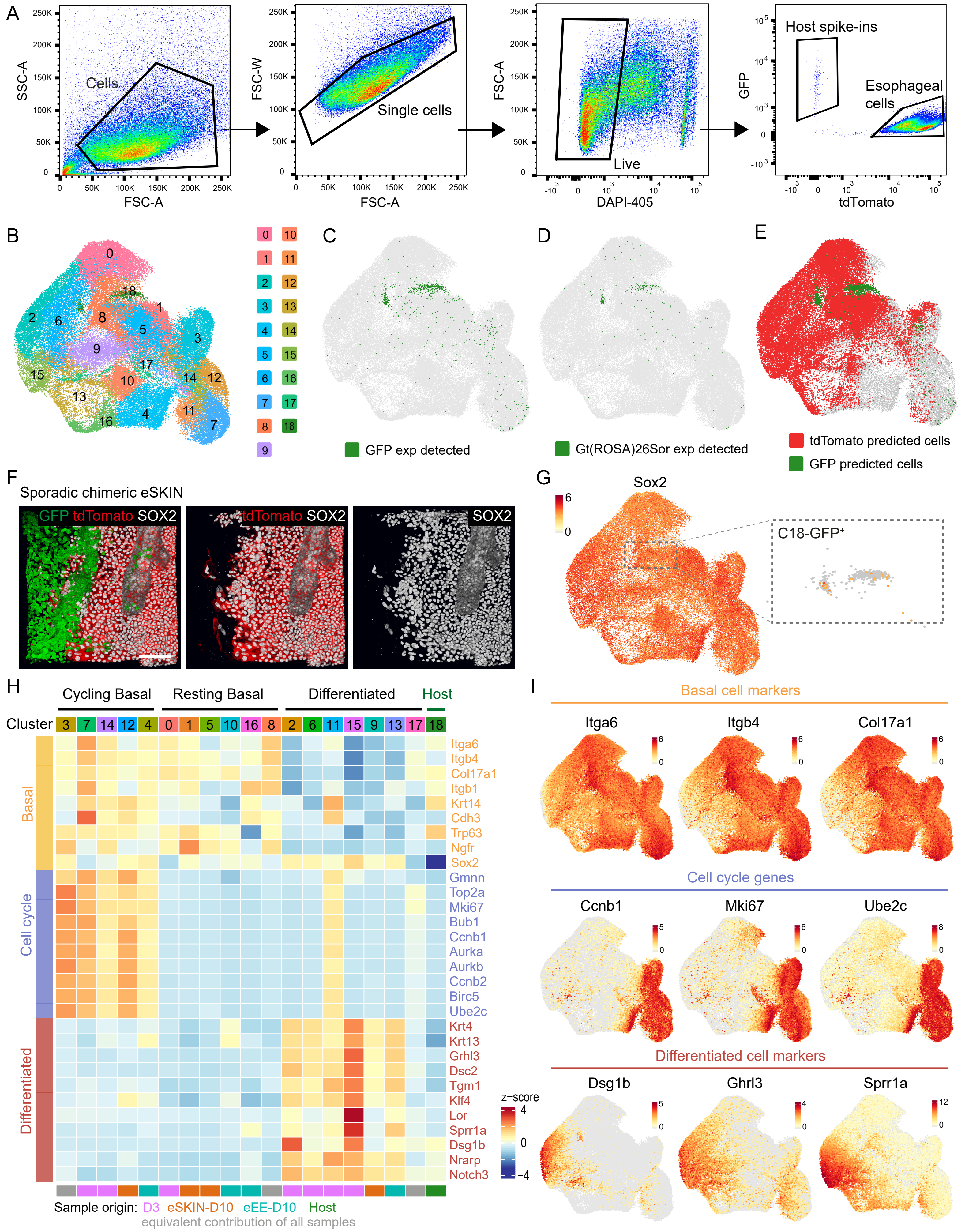
A
SSC-A
FSC-A
Cells
FSC-W
Single cells
Live
DAPI-405
GFP
Host spike-ins
Esophageal cells
tdTomato
B
C
GFP exp detected
D
Gt(ROSA)26Sor exp detected
E
tdTomato predicted cells
GFP predicted cells
F
Sporadic chimeric eSKIN
GFP tdTomato SOX2
tdTomato SOX2
SOX2
G
Sox2
C18-GFP+
H
Cycling Basal
Resting Basal
Differentiated
Host
Cluster
3 7 14 12 4 0 1 5 10 16 8 2 6 11 15 9 13 17 18
Basal
Cell cycle
Differentiated
Itga6
Itgb4
Col17a1
Itgb1
Krt14
Cdh3
Trp63
Ngfr
Sox2
Gmnn
Top2a
Mki67
Bub1
Ccnb1
Aurka
Aurkb
Ccnb2
Birc5
Ube2c
Krt4
Krt13
Grhl3
Dsc2
Tgm1
Klf4
Lor
Sprr1a
Dsg1b
Nrarp
Notch3
z-score
Sample origin: D3 eSKIN-D10 eEE-D10 Host
equivalent contribution of all samples
I
Basal cell markers
Itga6
Itgb4
Col17a1
Cell cycle genes
Ccnb1
Mki67
Ube2c
Differentiated cell markers
Dsg1b
Ghrl3
Sprr1a

**Supplementary Figure 3. Single-cell RNA sequencing annotation. Related to Figure 3. (A)** Flow cytometry gating strategy for isolation of esophageal (tdTomato) and host (EGFP) cells from 3D heterotypic cultures. Representative image from esophageal-derived skin (eSKIN) sample. 3-5 samples per time point were analyzed. **(B)** UMAP representing the cell allocation as a result of unsupervised clustering. **(C)** UMAP illustrating cells with detected expression of one of the segments of the EGFP sequence in C18 (host skin origin). **(D)** UMAP showing cells with detected expression of Gt(ROSA)26Sor -gene disrupted in esophageal cells (Rosa26$^{nTnG}$) due to insertion of the nTnG cassette. **(E)** UMAP representing prediction of cells carrying fluorescent labels tdTomato (EE) and EGFP (skin origin) based on a semi-supervised classifier that extends transcriptomic signatures of the detected tdTomato and EGFP cells to unlabeled (fluorescent marker genes not detected) cells. **(F)** Confocal images showing sporadic chimeric esophageal derived skin from heterotypic cultures tdTomato (nTnG; EE) and EGFP (skin origin). Note host cells are negative for typical EE Sox2 staining. Scale bar 50µm. **(G)** UMAP showing largely absent Sox2 expression in cluster 18. Inset represents magnifications of cells in cluster 18. **(H)** Heatmap showing expression of representative marker genes for basal cells, cell cycle, and differentiation for the clusters shown in **(B)**. **(I)** UMAPs showing expression of representative markers for basal (top panel), cell cycle (middle panel) and differentiated cells (bottom panel).

### Figure S4

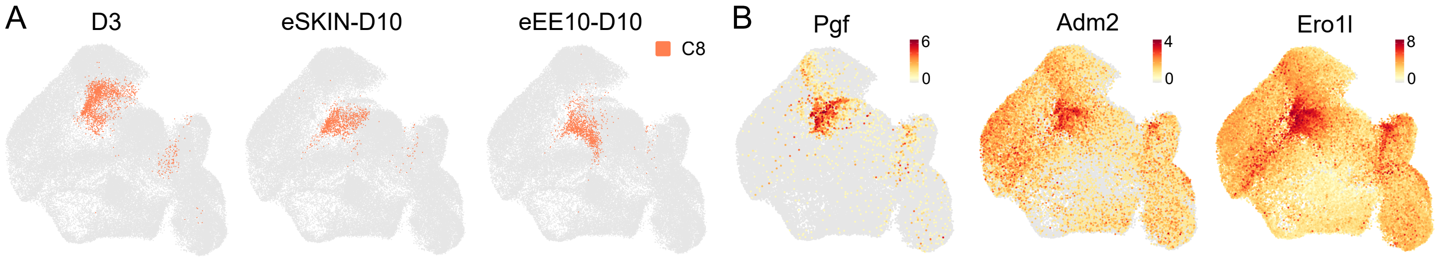


C

INCREASED DURING LINEAGE TRANSITION

| Pattern | P-value | Term | Genes |
|---|---|---|---|
| Pattern 3 | **Signaling and cellular response to stress** | | |
| | 2.11E-11<br>4.45E-05 | response to hypoxia [GO:BP]<br>response to stress [GO:BP] | Adm, Bnip3, Ddit4,<br>Ero1l, Ndrg1, Ptges |
| | 9.70E-08<br>3.47E-02 | HIF-1 signaling pathway [KEGG]<br>signaling receptor binding [GO:BP] | Egln1, Egln3, Hbegf,<br>Map2k1, Pgk1, Vegfa |
| | **Development and ECM remodeling** | | |
| | 2.83E-02<br>3.07E-02 | anatomical structure dev [GO:BP]<br>tube development [GO:BP] | Bsg, Cited1, Fstl3,<br>Pthlh, Rhob, Serpine1 |
| | 1.17E-02<br>1.56E-02 | Gap junction [KEGG]<br>ECM organization [REAC] | Adam8, Gja1, Itgb1,<br>Plod2, Tubb5, Tubb6 |
| | **Metabolism** | | |
| | 7.09E-06 | Glycolysis / Gluconeogenesis [KEGG] | Aldoc, Gapdh, Gpi1,<br>Pgam1, Pkm, Tpi1 |
| Pattern 2 | **Signaling and cellular response to stress** | | |
| | 9.29E-03 | cell population proliferation [GO:BP] | Cav1, Eif2s2, Fgfbp1,<br>Ndrg2, Pdgfa |
| | 1.34E-02 | response to stimulus [GO:BP] | Aqp3, Cst3, Fhl2,<br>Ier5, Klk8, Slc2a1 |
| | **Migration and ECM remodeling** | | |
| | 8.11E-02 | epithelial cell migration [GO:BP] | Apoe, Cd63, Klf4,<br>Mmp9, Sparc |

DECREASED DURING TRANSITION

| Pattern | P-value | Term | Genes |
|---|---|---|---|
| Pattern 1 | **Development and tissue differentiation** | | |
| | 7.47E-05 | anatomical structure development [GO:BP] | Dst, Wnt10a, Thbs1,<br>Id1, Wnt4, Hspd1 |
| | **Metabolism and translation** | | |
| | 2.97E-03<br>3.80E-02 | Ribosome biogenesis [KEGG]<br>rRNA processing [REAC] | Ddx21, Gnl3, Ncl, Nhp2,<br>Nop56, Nop58, Rexo2 |
| | 1.72E-02 | Glutathione metabolism [KEGG] | Idh1, Mgst1,<br>Mgst2, Prdx6 |
| Pattern 4 | **Metabolism** | | |
| | 1.03E-05 | Glutathione metabolism [KEGG] | Gclm, Gsr, Gsta4,<br>Gpx2, Mgst3, Odc1 |

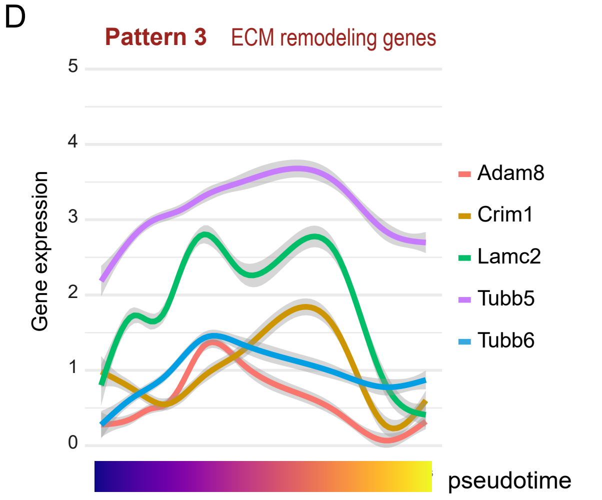


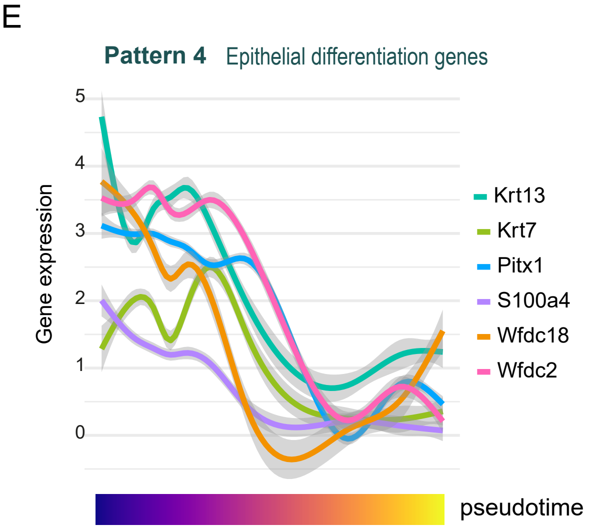

**Supplementary Figure 4. Gene expression patterns along the pseudotime trajectory that defines the identity transition. Related to Figure 4. (A)** UMAPs showing C8 in the three major transcriptional branches (**Fig3C**; D3, eEE-D10 and eSKIN-D10). **(B)** UMAPs showing maker genes of C8. **(C)** Table summarizing selected GO terms, KEGG, and REAC pathways for patterns 1-4, corresponding p-values and representative genes of patterns along the pseudotime trajectory. Related functional signatures are grouped together. **(D, E)** Expression of relevant genes along the pseudotime trajectory from esophageal to skin identity for ECM remodelling genes in pattern 3 **(D)** and epithelial differentiation genes in pattern 4 **(E)**.
